# Sclerostin blockade inhibits bone resorption through PDGF receptor signaling in osteoblast lineage cells

**DOI:** 10.1101/2023.09.11.557168

**Authors:** Cyril Thouverey, Pierre Apostolides, Julia Brun, Joseph Caverzasio, Serge Ferrari

## Abstract

While sclerostin-neutralizing antibodies (Scl-Ab) transiently stimulate bone formation by activating Wnt signaling in osteoblast lineage cells, they exert sustained inhibition of bone resorption, suggesting an alternate signaling pathway by which Scl-Ab control osteoclast activity. Since sclerostin can activate platelet-derived growth factor receptors (PDGFRs) in osteoblast lineage cells in vitro and PDGFR signaling in these cells induces bone resorption through M-CSF secretion, we hypothesized that the prolonged anti-catabolic effect of Scl-Ab could result from PDGFR inhibition. We show here that inhibition of PDGFR signaling in osteoblast lineage cells is sufficient and necessary to mediate prolonged Scl-Ab effect on M-CSF secretion and osteoclast activity in mice. Indeed, sclerostin co-activates PDGFRs independently of Wnt/β-catenin signaling inhibition, by forming a ternary complex with LRP6 and PDGFRs in pre-osteoblasts. In turn, Scl-Ab prevents sclerostin-mediated co-activation of PDGFR signaling and consequent M-CSF up-regulation in pre-osteoblast cultures, thereby inhibiting osteoclast activity in pre-osteoblast/osteoclast co-culture assays. These results provide a new potential mechanism explaining the dissociation between anabolic and anti-resorptive effects of long-term Scl-Ab.

## Introduction

Sclerostin, encoded by the *SOST* gene, is an osteocyte-secreted protein that antagonizes low-density lipoprotein receptor-related proteins (LRP) 5 and 6, thereby inhibiting canonical Wnt signaling in osteoblast lineage cells and bone formation (1–4). Due to its restricted expression in the adult skeleton, sclerostin has emerged as an attractive therapeutic target to increase bone mass and strength in osteoporotic patients. Consequently, administration of antibodies targeting sclerostin (Scl-Ab) has been shown to augment bone mineral density and bone strength in humans, through transient elevation of bone formation and sustained reduction of bone resorption (5–10).

Mechanistically, short-term Scl-Ab treatment induces a rapid and intense increases in serum procollagen type I N-terminal propeptide (PINP) and histomorphometric indices of bone formation, as well as a significant decrease in serum level of the bone resorption marker C-terminal telopeptides of type I collagen (CTX) in osteoporotic patients (7–10). Preclinical investigations have shown that Scl-Ab simultaneously activate modeling-based bone formation by stimulating transition of bone lining cells into active osteoblasts and generate a positive bone balance at remodeling sites by enhancing the anabolic power (vigor) of each osteoblast (11–13). In parallel, bone resorption surfaces are decreased and associated with lower receptor activator of nuclear factor κB ligand (RANKL)/osteoprotegerin (OPG) ratio, decreased expression of *Csf1* (encoding macrophage colony-stimulating factor, M-CSF), essential for osteoclast differentiation and survival, and enhanced expression of *Wisp1*, a negative regulator of bone resorption (13, 14).

After 3 to 6 months of Scl-Ab treatment, serum PINP and bone formation indices return to initial values, whereas serum CTX and bone resorption parameters are maintained below baseline levels (7–10). Although overall bone turnover eventually decreases, the positive bone mineral balance within bone remodeling units is maintained, but attenuated, allowing significant bone mass gain to continue for the duration of therapy (1 year) (15). Counter-regulation of bone formation with Scl-Ab administration can be explained by increased expressions of Wnt pathway inhibitors such as *Sost* and *Dkk1* (encoding dickkopf-related protein 1, DKK1), and a decreasing number of osteoprogenitors (14, 16). In this context, the reason why Scl-Ab exerts prolonged anti-catabolic effect despite attenuation of its bone anabolic activity remains unclear. A possible explanation is that sclerostin neutralization could reduce bone resorption independently of canonical Wnt signaling activation.

We previously showed that platelet-derived growth factor receptor (PDGFR)α and PDGFRβ in *Osterix*-positive cells redundantly control osteoclastogenesis and bone resorption by upregulating expression of *Csf1* in mice (17), and that sclerostin can induce PDGFR signaling in osteoblast lineage cells in vitro (18). Therefore, we hypothesized that the prolonged anti-catabolic effect of Scl-Ab treatment could result from an inhibition of PDGFR signaling independently of its Wnt-activating properties in osteoblast lineage cells.

## Results

### Scl-Ab transiently stimulates bone formation in both control and *Pdgfr cKO* mice

To test the role of PDGFR signaling in osteoblast lineage cells on Scl-Ab effects in vivo, we treated *Osx-Cre;Pdgfra^f/f^;Pdgfrb^f/f^* (hereafter *Pdgfr cKO*) and control (*Osx-Cre*) mice with Scl-Ab or its vehicle solution (Veh) for 2 and 6 weeks (Figure 1A). Due to the short duration of PDGFR deletion (inducible Cre activation one week prior to treatment) (Figure 1B), *Pdgfr cKO* and *Osx-Cre* mice treated with Veh showed similar cortical and trabecular bone mass at all time points (Figure 1, C and D). Scl-Ab increased cortical bone volume at tibial midshaft, as well as trabecular number, thickness and bone volume at the proximal metaphysis at 2 and 6 weeks in both genotypes (Figure 1, C and D, and Supplemental Figure 1A). However, Scl-Ab increased trabecular bone volume more in *Pdgfr cKO* mice than in *Osx-Cre* mice after 2 weeks, a trend which persisted after 6 weeks of treatment (Figure 1D and Supplemental Figure 1A). Scl-Ab treatment initially stimulated the mineral apposition rate on trabecular bone surfaces equally in both genotypes (i.e. at 2 weeks), but increased trabecular mineralizing surfaces and bone formation rate more in *Pdgfr cKO* mice than in *Osx-Cre* mice (Figure 1E and Supplemental Figure 1, B and C). Bone formation parameters returned to pre-treatment levels after 6 weeks of Scl-Ab treatment in mice of both genotypes (Figure 1E and Supplemental Figure 1, B and C). Similarly, serum levels of type I procollagen N-terminal propeptide (PINP) tended to peak at a higher level in *Pdgfr cKO* mice than in *Osx-Cre* mice after 2 weeks of Scl-Ab treatment, but declined to similar levels in both genotypes after 6 weeks of treatment (Figure 1F). These observations indicated that PDGFR signaling could attenuate the extent of bone-forming surfaces in response to Scl-Ab, but did not play a role in the counter-regulation of Scl-Ab effect on bone formation.

**Figure 1.**
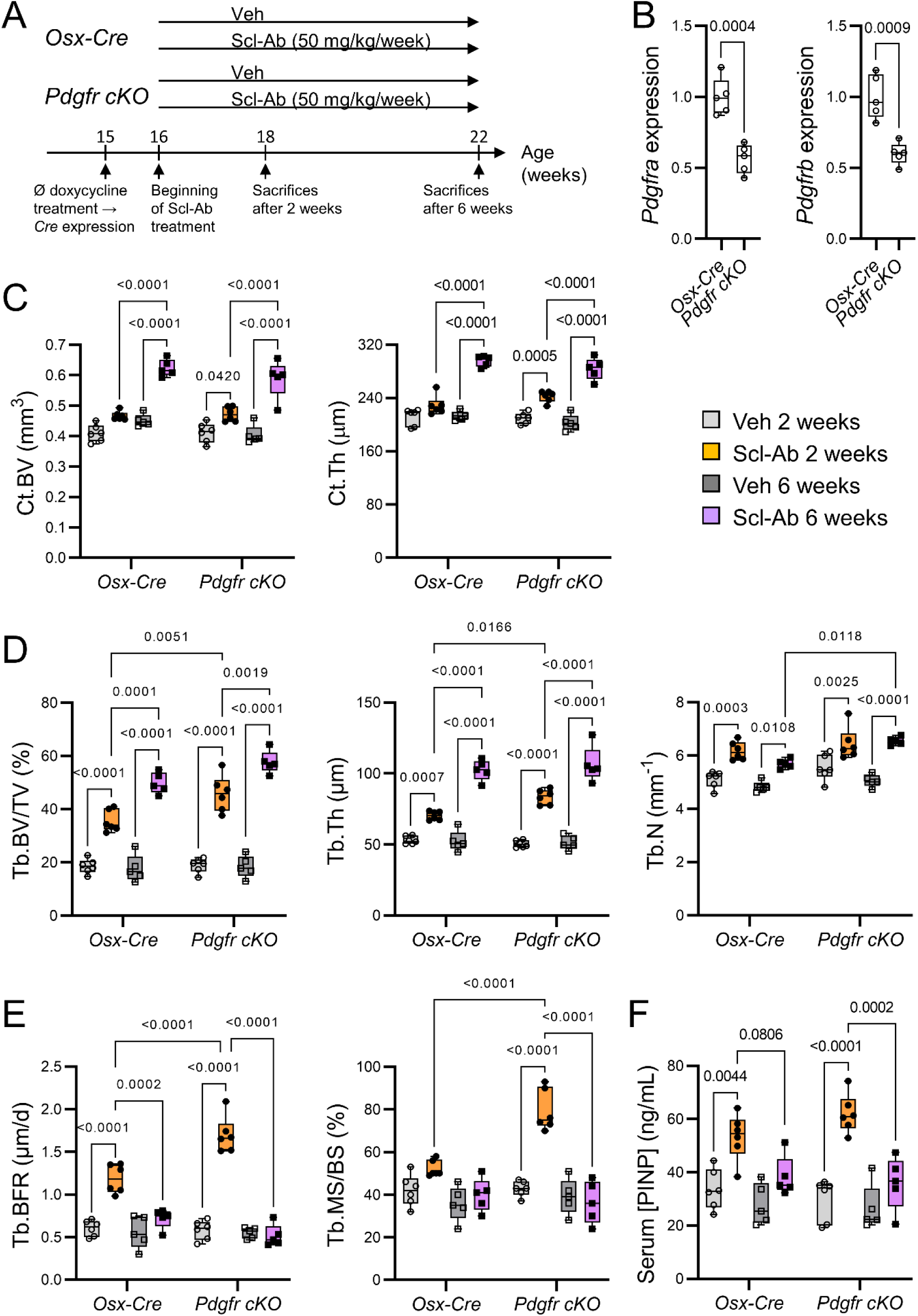
Scl-Ab-induced bone formation peaked more strongly in *Pdgfr cKO* mice than in control mice after two weeks of treatment, but declined to baseline value after six weeks of Scl-Ab treatment in mice of both genotypes. (A) 4-month-old *Osx-Cre* and *Pdgfr cKO* (*Pdgfra cKO;Pdgfrb cKO*) male mice received subcutaneous injections of saline solution (Veh) or 25 mg/kg Scl-Ab twice a week for 2 or 6 weeks. *Cre* expression or/and conditional gene deletion were induced (by stopping doxycycline treatment) one week prior the beginning of Scl-Ab treatment. (B) Quantitative RT-PCR analyses of *Pdgfra* and *Pdgfrb* expressions in bone marrow-free proximal tibial metaphyses at baseline (n=5 per group). Differences between the two genotypes were analyzed using unpaired t-tests. (C-E) Concerning µCT, histomorphometric analyses and serum levels of PINP, interactions between effects of genotypes, those of treatments, and those of treatment durations were analyzed by linear mixed-effects models. Differences between genotypes, between treatments or between durations were analyzed using Tukey post hoc tests. (C) Cortical bone volume (Ct.BV) and thickness (Ct.Th) measured at tibial midshaft (n=5-6 per group). (D) Trabecular bone microarchitecture measured at proximal tibiae (n=5-6 per group). µCT parameters include BV/TV: bone volume/total volume; Tb.Th: trabecular thickness; Tb.N: trabecular number. (E) Histomorphometric parameters of trabecular bone formation measured at the secondary spongiosa of distal femurs (n=5-6 per group). Tb.BFR: trabecular bone formation rate; Tb.MS/BS: trabecular mineralizing surfaces/bone surfaces. (F) Serum levels of PINP (n=5-6 per group).

### Self-regulation of Wnt signaling in response to Scl-Ab is independent of PDGFRs

The similar decline of Scl-Ab anabolic effects after 6 weeks in both *Pdgfr cKO* and control mice suggested that prolonged Scl-Ab treatment is associated with a self-regulation of Wnt signaling that is independent of PDGFR signaling. Hence, we measured expressions of selected Wnt target genes and Wnt pathway regulators in proximal tibial metaphysis isolated from *Osx-Cre* and *Pdgfr cKO* mice. 2-week Scl-Ab treatment enhanced expressions of *Wisp1* and *Twist1*, two Wnt target genes which are responsive to sclerostin neutralization (13, 14), even more strongly in *Pdgfr cKO* mice than in *Osx-Cre* mice (Figure 2, A and B). Expressions of both Wnt target genes returned to basal levels following 6 weeks of Scl-Ab treatment in both genotypes, thereby confirming attenuation of Wnt signaling with prolonged exposure to Scl-Ab (Figure 2, A and B). This down-regulation of Wnt target gene expression was associated with a significant elevation of *Sost* (after 6 weeks) and *Dkk1* (from 2 weeks) expressions in both genotypes (Figure 2, C and D). Moreover, Scl-Ab administration also quickly reduced expressions of *Wnt1* and *Wnt10b* in bones of mice from both genotypes (Figure 2, E and F).

**Figure 2.**
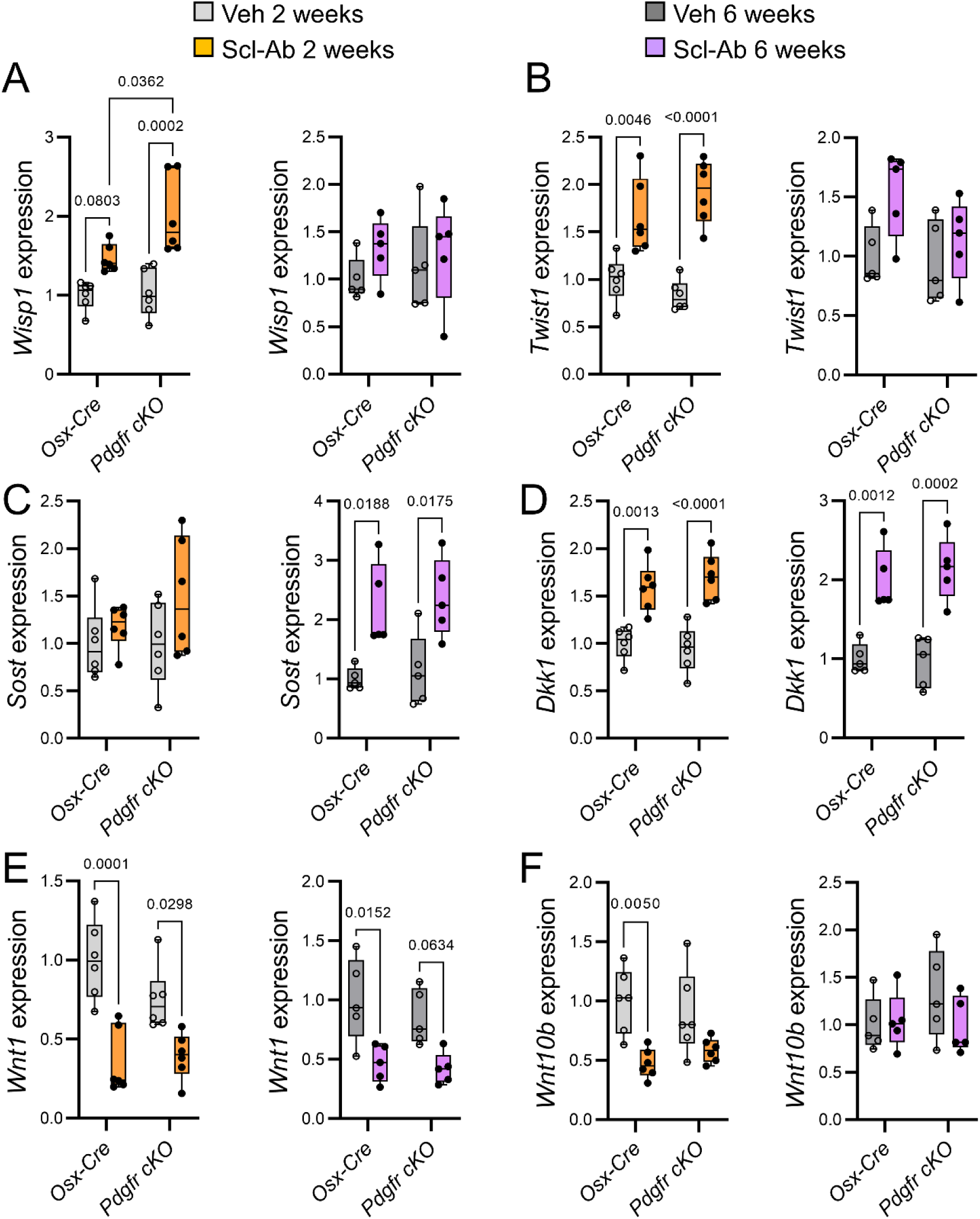
Self-regulation of Scl-Ab-induced Wnt signaling in bone coincided with increased expression of sclerostin and decreased expressions of Wnt-1 class of proteins in mice of both genotypes. 4-month-old *Osx-Cre* and *Pdgfr cKO* male mice received subcutaneous injections of saline solution (Veh) or 25 mg/kg Scl-Ab twice a week for 2 weeks or 6 weeks. *Cre* expression or/and conditional gene deletion were induced one week prior the onset of Scl-Ab treatment. Interactions between effects of genotypes and those of treatments were analyzed by 2-way ANOVA. Differences between genotypes or between treatments were performed using Tukey post hoc tests. (A-F) Quantitative RT-PCR analyses of (A) *Wisp1* (encoding Wnt1-inducible-signaling pathway protein 1), (B) *Twist1* (twist-related protein 1), (C) *Sost* (sclerostin), (D) *Dkk1* (Dickkopf-related protein 1), (E) *Wnt1* and (F) *Wnt10b* expressions in proximal tibial metaphyses (n=5-6 per group).

Altogether, those results indicated that Wnt signaling attenuation following prolonged Scl-Ab treatment was mediated by a negative feedback mechanism involving increased expressions of Wnt signaling inhibitors and decreased expressions of Wnt1 class of ligands, independently of PDGFR signaling.

### PDGFRs mediate Scl-Ab inhibitory effects on bone resorption and M-CSF secretion

The suppression of PDGFRs in osteoblast lineage cells significantly decreased osteoclast number and surfaces and tended to reduce serum tartrate-resistant acid phosphatase isoform 5b (TRAcP 5b) levels (Figure 3, A-D), together with reduced *Csf1* expression and M-CSF protein levels in the bone micro-environment (Figure 3, E and F). Scl-Ab significantly reduced osteoclast number and surfaces in control mice after 2 and 6 weeks (Figure 3, A-C), while it tended to further diminish the already low osteoclast number and surfaces in *Pdgfr cKO* mice after 2 weeks, but not after 6 weeks of treatment (Figure 3, A-C). Scl-Ab also tended to decrease serum levels of TRAcP 5b in *Osx-Cre* mice, but not in *Pdgfr cKO* mice (Figure 3D). Accordingly, sclerostin neutralization lowered *Csf1* expression in bone and M-CSF protein levels in the bone marrow of control *Osx-Cre* mice but not in those of *Pdgfr cKO* mice after 2 and 6 weeks of treatment (Figure 3, E and F). In contrast, we found no significant differences in *Rankl* or *Opg* expression in response to Scl-Ab treatment and PDGFRs deletion (Figure 3, G and H).

**Figure 3.**
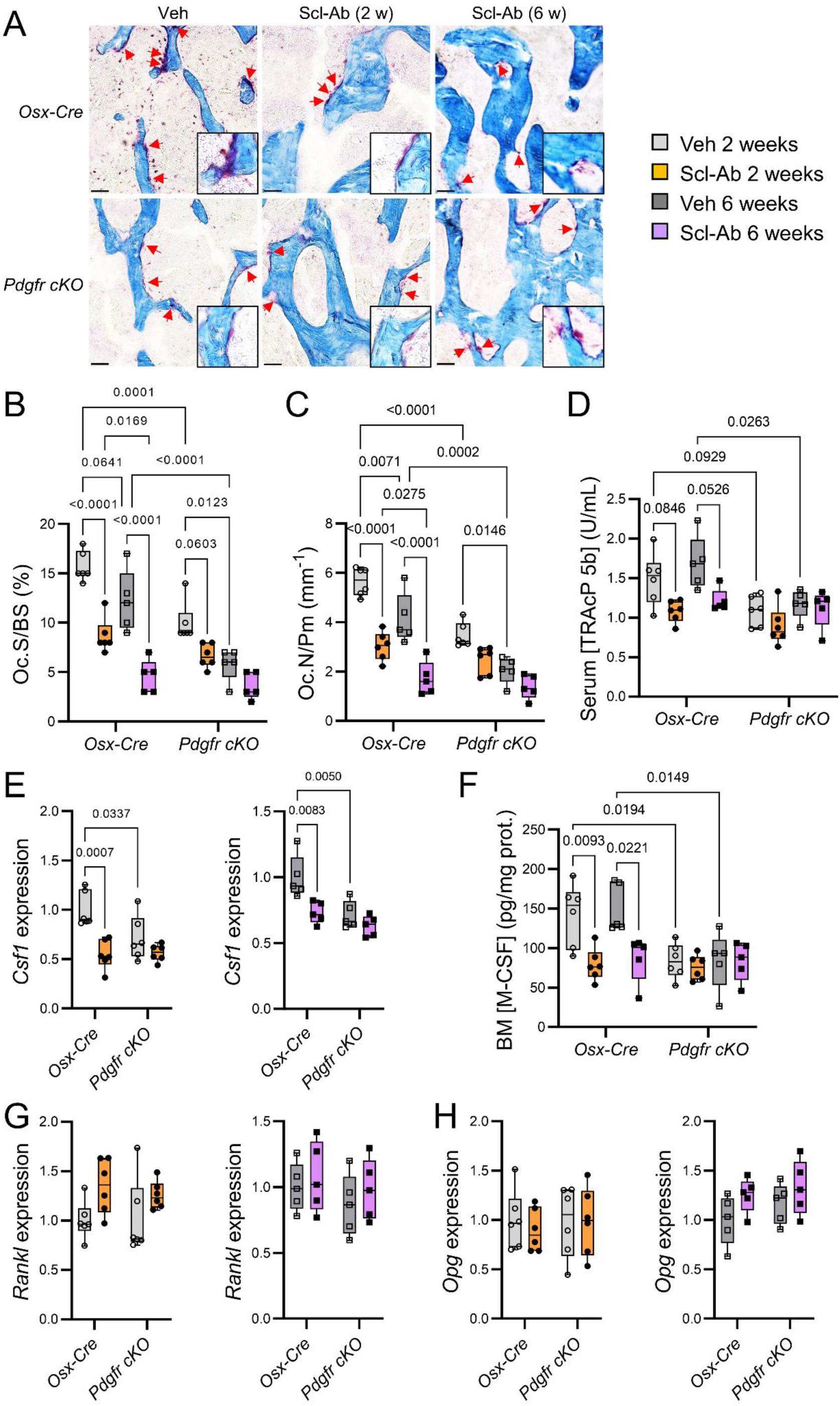
Scl-Ab treatment durably decreased bone resorption and *Csf1* expression in control mice, but did not exert any further anti-catabolic effect in *Pdgfr cKO* mice. 4-month-old *Osx-Cre* and *Pdgfr cKO* (*Pdgfra cKO;Pdgfrb cKO*) male mice received subcutaneous injections of saline solution (Veh) or 25 mg/kg Scl-Ab twice a week for 2 or 6 weeks. *Cre* expression or/and conditional gene deletion were induced (by stopping doxycycline treatment) one week prior the beginning of Scl-Ab treatment. Interactions between effects of genotypes, those of treatments, and those of treatment durations were analyzed by linear mixed-effects models. Differences between genotypes, between treatments or between durations were analyzed using Tukey post hoc tests. (A) Representative images of TRAP-stained histological sections of distal femurs (Scale bars represent 50 µm). (B, C) Histomorphometric parameters of trabecular bone resorption measured at the secondary spongiosa of distal femurs (n=5-6 per group). Oc.S/BS: osteoclast surface/bone surface; Oc.N/Pm: osteoclast number/bone perimeter. (D) Serum levels of TRAcP 5b (n=5-6 per group). (E) Quantitative RT-PCR analyses of *Csf1* (encoding M-CSF) expression in proximal tibial metaphyses (n=5-6 per group). (F) M-CSF protein levels in bone marrow (BM) supernatants (n=5-6 per group). (G, H) Quantitative RT-PCR analyses of (G) *Rankl* (receptor activator of nuclear factor κB ligand) and (H) *Opg* (osteoprotegerin) expressions in proximal tibial metaphyses (n=5-6 per group).

Those data indicated that sclerostin and PDGFRs operated within the same signaling pathway to durably inhibit M-CSF secretion and bone resorption.

### Scl-Ab inhibits expression of PDGFR-responsive genes in bone without down-regulating PDGFRs expression

To confirm that Scl-Ab-mediated inhibition of bone resorption is associated with a concomitant inhibition of PDGFR signaling, we measured expressions of PDGFR target genes and PDGFR signaling components in proximal tibial metaphysis isolated from *Osx-Cre* and *Pdgfr cKO* mice treated with Veh and Scl-Ab for 2 and 6 weeks. Selective suppressions of *Pdgfra* and *Pdgfrb* in osteoblast lineage cells were associated with reduced expressions of *c-Myc* and *Ccl2* (Figure 4, A and B), two PDGFR target genes (19, 20), and confirmed by reduced expressions of both genes in *Pdgfr cKO* mice (Figure 4, C and D). In contrast to the transient effect of Scl-Ab on Wnt target gene expression (Figure 2), Scl-Ab treatment consistently decreased expressions of *c-Myc* and *Ccl2* in *Osx-Cre* mice consistently after 2 and 6 weeks, but did not further reduce the already low expression levels of these genes in *Pdgfr cKO* mice (Figure 4, A and B). In contrast, Scl-Ab did not affect *Pdgfra* and *Pdgfrb* expression levels (Figure 4, C and D). Furthermore, both selective PDGFRs ablation in osteoblast lineage cells and Scl-Ab administration reduced expression of *Pdgfb*, encoding PDGF-B ligand (Figure 4E), probably reflecting low number of *Pdgfb*-expressing osteoclasts under these conditions (17). These data indicated that Scl-Ab directly inhibited the expression of PDGFR signaling target genes and these effects were not counter-regulated by the increased expression of Wnt inhibitors.

**Figure 4.**
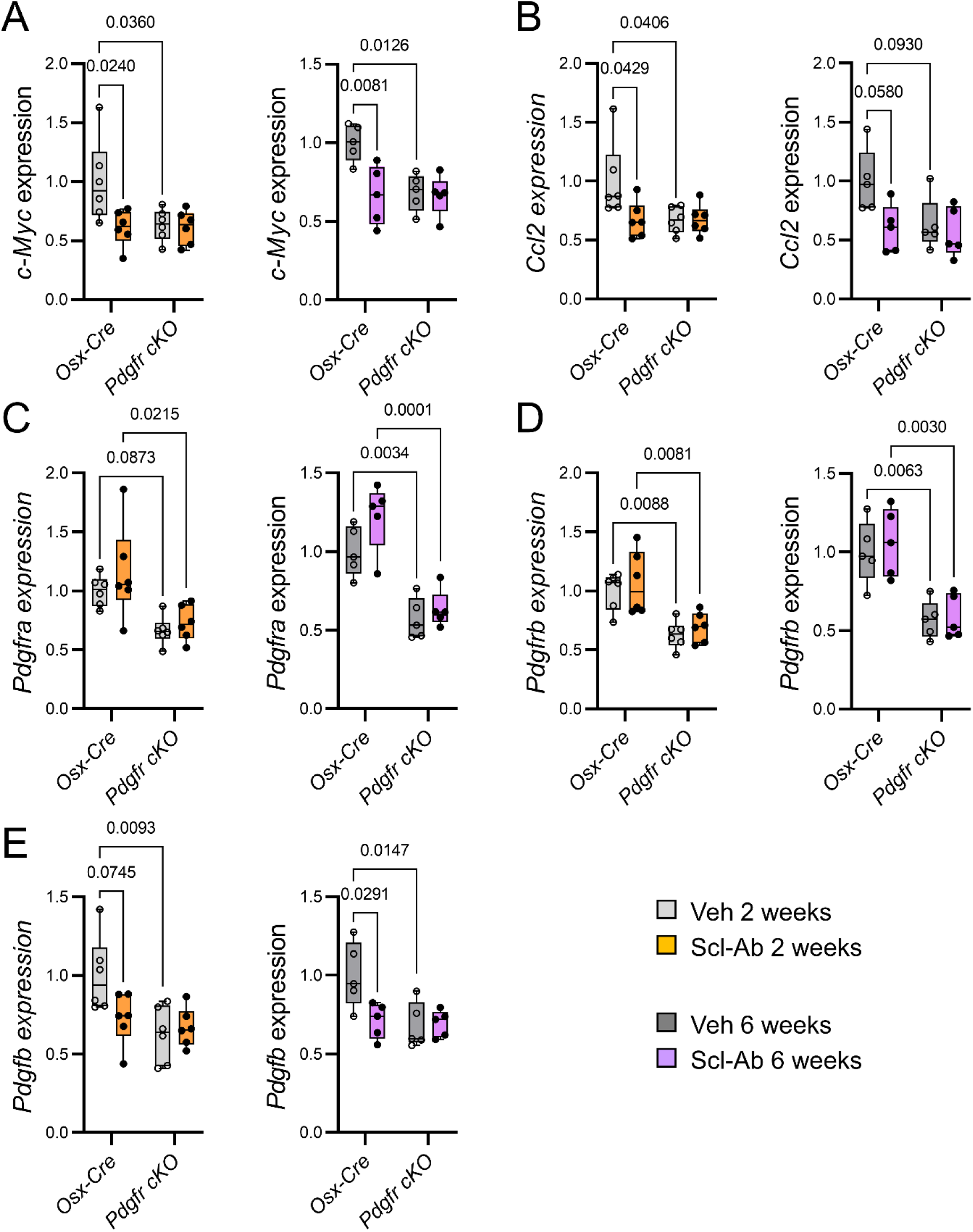
Scl-Ab treatment durably decreased expressions of PDGFR target genes in bone of control mice, but did not exert any further inhibitory effect in *Pdgfr cKO* mice. 4-month-old *Osx-Cre* and *Pdgfr cKO* male mice received subcutaneous injections of saline solution (Veh) or 25 mg/kg Scl-Ab twice a week for 2 weeks or 6 weeks. *Cre* expression or/and conditional gene deletion were induced one week prior the onset of Scl-Ab treatment. Interactions between effects of genotypes and those of treatments were analyzed by 2-way ANOVA. Differences between genotypes or between treatments were performed using Tukey post hoc tests. (A-E) Quantitative RT-PCR analyses of (A) *c-Myc* (encoding myc pro-oncogen protein), (B) *Ccl2* (C-C motif chemokine 2), (C) *Pdgfra* (PDGFRα), (D) *Pdgfrb* (PDGFRβ) and (E) *Pdgfb* (PDGF-B) expressions in proximal tibial metaphyses (n=5-6 per group).

### Sclerostin potentiates PDGF-BB-stimulated *Csf1* expression and bone resorption in vitro by forming a sclerostin/LRP6/PDGFR ternary complex for ERK1/2 activation

To confirm and expand these in vivo observations, we analyzed the molecular mechanisms by which sclerostin could interfere with PDGFR signaling to regulate osteoclast development and function in vitro. Since PDGFRα and PDGFRβ expressions were high in early osteoblasts expressing *Runx2* and *Col1a1*, and then declined as those cells differentiated into mature osteoblasts expressing *Ocn*, we used primary cultures of pre-osteoblasts (Supplemental Figure 2). Recombinant sclerostin did not have any effect on in vitro osteoclastogenesis in basal conditions, but enhanced calcitriol-induced osteoclastogenesis and demineralization of a synthetic matrix in pre-osteoblast/osteoclast co-cultures (Figure 5, A and B, and Supplemental Figure 3, A and B). These effects were blocked by Scl-Ab, M-CSF-targeting antibody or by PDGFR deletion in pre-osteoblasts (Figure 5, A and B, and Supplemental Figure 3, A and B). In addition, sclerostin potentiated PDGF-BB-induced *Csf1* expression and M-CSF secretion in pre-osteoblast cultures (Figure 5, C and D). Again, those effects were blocked by either Scl-Ab treatment (Figure 5, C and D) or suppression of PGDFRs in pre-osteoblasts (Figure 5C). It should be noted that sclerostin, Scl-Ab or PDGFR deletion did not alter osteoblast differentiation during those cell culture experiments (Supplemental Figure 3, B and C). Consistent with those findings, sclerostin amplified PDGF-BB-induced phosphorylation of PDGFRs in pre-osteoblast cultures after 15 minutes and elevation of M-CSF protein level after 24 hours, effects that were abrogated in the presence of Scl-Ab (Figure 5E). Eventually, immunoprecipitation of proteins by anti-LRP6 antibody showed that sclerostin could intensify PDGF-BB-induced activation of PDGFRs by forming a ternary complex with LRP6 and PDGFRs in pre-osteoblasts (Figure 5F). Sclerostin-mediated potentiating effects on PDGF-BB-dependent PDGFRs activation resulted in further stimulation of downstream PDGFR signaling pathways such as ERK1/2 and STAT3 in pre-osteoblasts (Figure 5G).

**Figure 5.**
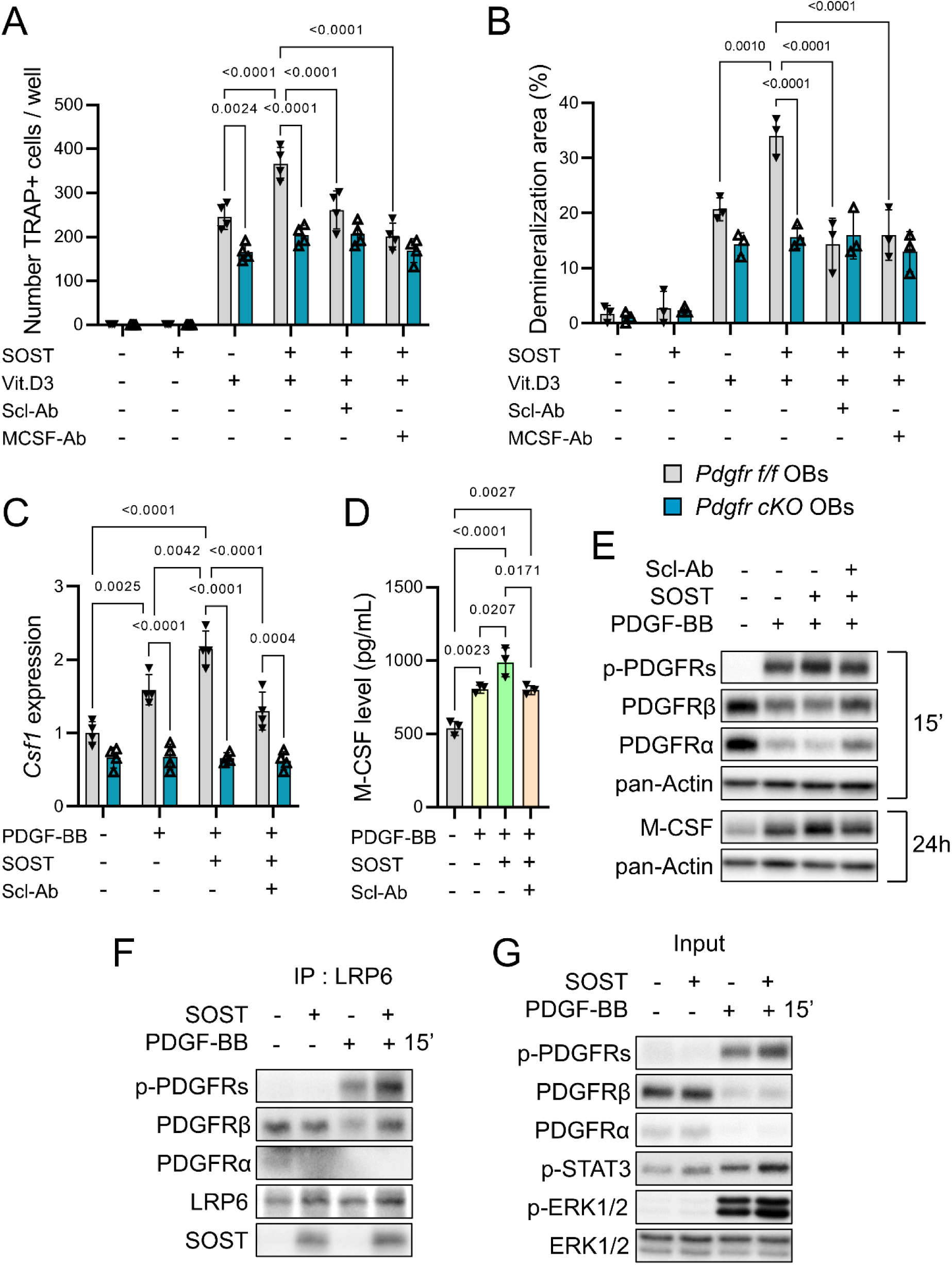
Scl-Ab blocked sclerostin-mediated PDGFR co-activation and M-CSF secretion in pre-osteoblast cultures, and calcitriol-induced osteoclast formation and activity in co-cultures. (A) *Pdgfr^f/f^* (control) and *Pdgfr cKO* (without PDGFRs) pre-osteoblasts were co-cultured with non-adherent bone marrow cells isolated from wildtype mice in the presence of Veh, 10^-8^ M 1,25-dihydroxyvitamin D3 (calcitriol, Vit.D3), or/and 250 ng/mL recombinant sclerostin (SOST), with or without 1.25 µg/mL Scl-Ab or 500 ng/mL anti-M-CSF antibody (MCSF-Ab) during 8 days before quantification of TRAP-positive multinucleated cells. (B) The same co-cultures performed in synthetic matrix-coated multiwell plates during 15 days before quantification of demineralized areas. (C) *Pdgfr^f/f^* and *Pdgfr cKO* pre-osteoblasts were pre-treated with Veh, 250 ng/mL SOST ± 1.25 µg/mL Scl-Ab for 1 hour, and treated with Veh or 25 ng/mL PDGF-BB for 24 hours before measurements of *Csf1* expression by quantitative RT-PCR. (D) Wildtype pre-osteoblasts were pre-treated with Veh, 250 ng/mL SOST ± 1.25 µg/mL Scl-Ab for 1 hour, and treated with Veh or 25 ng/mL PDGF-BB for 48 hours before quantification of M-CSF in culture media by ELISA tests. (E) Wildtype pre-osteoblasts were pre-treated with Veh or 500 ng/mL SOST ± 2.5 µg/mL Scl-Ab for 1 hour, and then treated with 15 ng/mL PDGF-BB for indicated time periods. PDGFR signaling and M-CSF protein level were determined by western blot analyses. (F, G) Wildtype pre-osteoblasts were pre-treated with Veh or 500 ng/mL SOST for 2 hours, and then treated with 25 ng/mL PDGF-BB for 15 minutes. (F) Proteins were immunoprecipitated by anti-LRP6 antibody and detected by western blot analyses. (G) PDGFR signaling was assessed in the remaining cell lysates.

### Sclerostin potentiates PDGF-BB-induced *Csf1* expression independently of Wnt/β-catenin signaling in pre-osteoblast cultures

To provide an explanation for the continuous inhibition of PDGFR signaling and *Csf1* expression despite self-attenuation of Wnt signaling in response to prolonged Scl-Ab exposure (Figure 4), we tested whether Wnt/β-catenin signaling activation or inhibition could influence sclerostin ability to co-activate PDGFRs and stimulate *Csf1* expression in pre-osteoblast cultures. First, the presence of Wnt1 in culture medium did not alter potentiating effects of sclerostin on PDGF-BB-induced *Csf1* expression, M-CSF production and PDGFR signaling activation (Figure 6, A-C). Second, inhibition of β-catenin signaling by WIKI4 had no effect on PDGF-BB-dependent sclerostin-mediated *Csf1* expression (Figure 6D). Third, in contrast to sclerostin, another Wnt signaling inhibitor, DKK1, could not potentiate PDGF-BB-mediated *Csf1* expression and PDGFRs/ERK1/2 activation (Figure 6, E and F). Together, those results indicated that sclerostin could potentiate PDGF-BB-mediated *Csf1* expression and PDGFRs activation independently of Wnt/β-catenin signaling inhibition. As a control, sclerostin capability to inhibit Wnt1-induced β-catenin activation and *Wisp1* expression was confirmed in pre-osteoblast cultures (Figure 6, G and H). In this context, it is noteworthy that suppression of PDGFRs in pre-osteoblasts could potentiate Wnt1-promoted β-catenin activation and *Wisp1* expression (Figure 6, G and H), thereby also explaining potentiating effect of osteoblast lineage-selective ablation of PDGFRs on Scl-Ab-stimulated bone formation (Figure 1).

**Figure 6.**
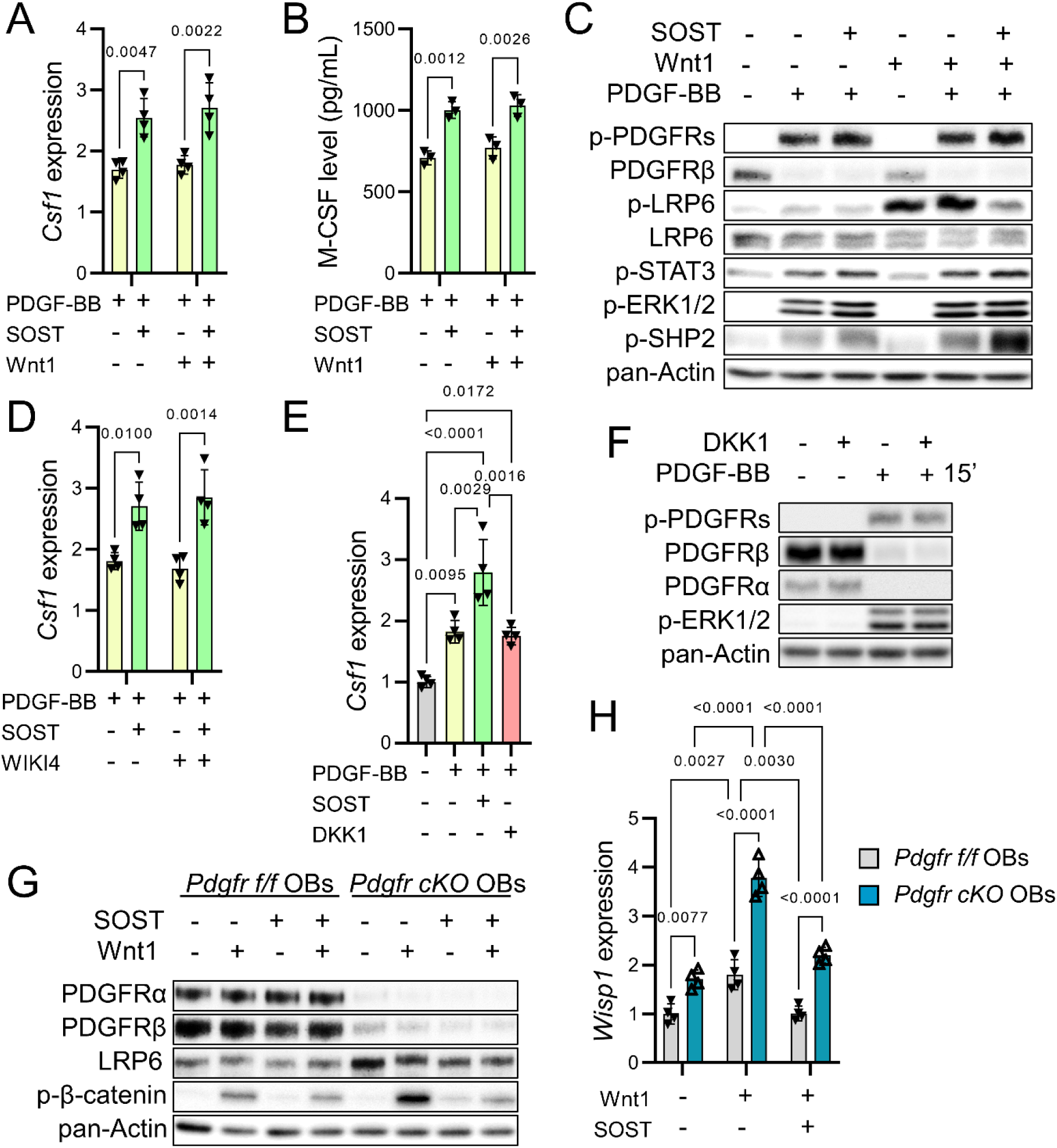
Sclerostin increases *Csf1* expression independently of Wnt/β-catenin signaling inhibition in pre-osteoblast cultures. (A, B) Wildtype pre-osteoblasts were pre-treated with Veh or 100 ng/mL Wnt1 ± 500 ng/mL SOST for 2 hours, and treated with Veh or 25 ng/mL PDGF-BB for (A) 24 hours before measurements of *Csf1* expression by quantitative RT-PCR, or (B) 48 hours before quantification of M-CSF in culture media by ELISA tests. (C) Wildtype pre-osteoblasts were pre-treated with Veh or 100 ng/mL Wnt1 ± 500 ng/mL SOST for 2 hours, and treated with Veh or 25 ng/mL PDGF-BB for 15 minutes before evaluation of PDGFR signaling by western blot analyses. (D) Wildtype pre-osteoblasts were pre-treated with DMSO or 5 µM WIKI4 (inhibitor of Wnt/β-catenin signaling), and Veh or 500 ng/mL SOST for 2 hours, and treated with Veh or 25 ng/mL PDGF-BB for 24 hours before measurements of *Csf1* expression by quantitative RT-PCR. (E) Wildtype pre-osteoblasts were pre-treated with Veh, 500 ng/mL SOST or 500 ng/mL DKK1 for 2 hours, and treated with Veh or 25 ng/mL PDGF-BB for 24 hours before measurements of *Csf1* expression by quantitative RT-PCR. (F) Wildtype pre-osteoblasts were pre-treated with Veh or 500 ng/mL DKK1 for 2 hours, and treated with Veh or 25 ng/mL PDGF-BB for 15 minutes before evaluation of PDGFR signaling by western blot analyses. (G) *Pdgfr^f/f^* and *Pdgfr cKO* pre-osteoblasts were pre-treated with Veh or 500 ng/mL SOST, and treated with Veh or 100 ng/mL Wnt1 for 2 hours before determination of Wnt/β-catenin signaling by western blot analyses. (H) *Pdgfr^f/f^* and *Pdgfr cKO* pre-osteoblasts were pre-treated with Veh or 500 ng/mL SOST, and treated with Veh or 100 ng/mL Wnt1 for 24 hours before measurements of *Wisp1* expression by quantitative RT-PCR.

## Discussion

Sclerostin blockade exerts dual effects on bone, resulting in a potent but transient stimulation of bone formation but a milder and sustained inhibition of bone resorption. Because Scl-Ab effects on bone formation are primarily mediated by stimulation of the Wnt/β-catenin signaling pathway in osteoblast lineage cells and this pathway is rapidly down-regulated by the overexpression of Wnt inhibitors including sclerostin itself and DKK1 (14, 16), the persistent inhibition of bone resorption suggests that sclerostin and its pharmacological inhibitors control osteoclastogenesis by an alternative, Wnt-independent pathway, in osteoblast lineage cells. Consistent with human data, we showed that, despite continuous cortical and trabecular bone mass gain, bone formation induced by Scl-Ab treatment in mice was transient. The decline of Scl-Ab osteoanabolic effects was associated with attenuation of Wnt signaling due to elevated expressions of Wnt signaling inhibitors, and decreased expressions of Wnt1 class of ligands. In contrast, bone PDGFR signaling, *Csf1* expression, M-CSF secretion and bone resorption were durably reduced by Scl-Ab in control mice. *Pdgfr cKO* mice recapitulated these changes and, although short-term Scl-Ab treatment tended to further decrease bone resorption in these mice, prolonged Scl-Ab exposure did not. Scl-Ab abolished sclerostin-mediated co-activation of PDGFR signaling and consequent M-CSF up-regulation in pre-osteoblast cultures, and osteoclast formation and activity in pre-osteoblast/osteoclast co-culture assays. Eventually, we showed that sclerostin could potentiate PDGFRs activation, unlike DKK1 and independently of the presence of Wnt ligand, by forming a ternary complex with LRP6 and PDGFRs in pre-osteoblasts.

Genetic ablation of PDGFRs-encoding genes potentiated Scl-Ab-promoted trabecular bone mass gain. This effect was mainly due to an amplified early bone anabolic response to sclerostin neutralization (Figure 1). From a mechanistic point of view, deletion of PDGFRs-encoding genes in pre-osteoblasts stimulated Wnt1-induced accumulation of active β-catenin and Wnt target gene expression (Figures 2 and 6), showing that PDGFRs negatively regulate the Wnt/β-catenin signaling pathway (Figure 7). The stimulation of bone formation by Scl-Ab therapy was only transient (Figure 1) (7–10, 14, 21), and the associated counter-regulation of canonical Wnt signaling occurred in both control and *Pdgfr cKO* mice (Figure 2), indicating that inhibitory action of PDGFRs on Wnt/β-catenin signaling in presence of Scl-Ab is overridden by DKK1 and low Wnt1 class availability (Figure 7).

**Figure 7.**
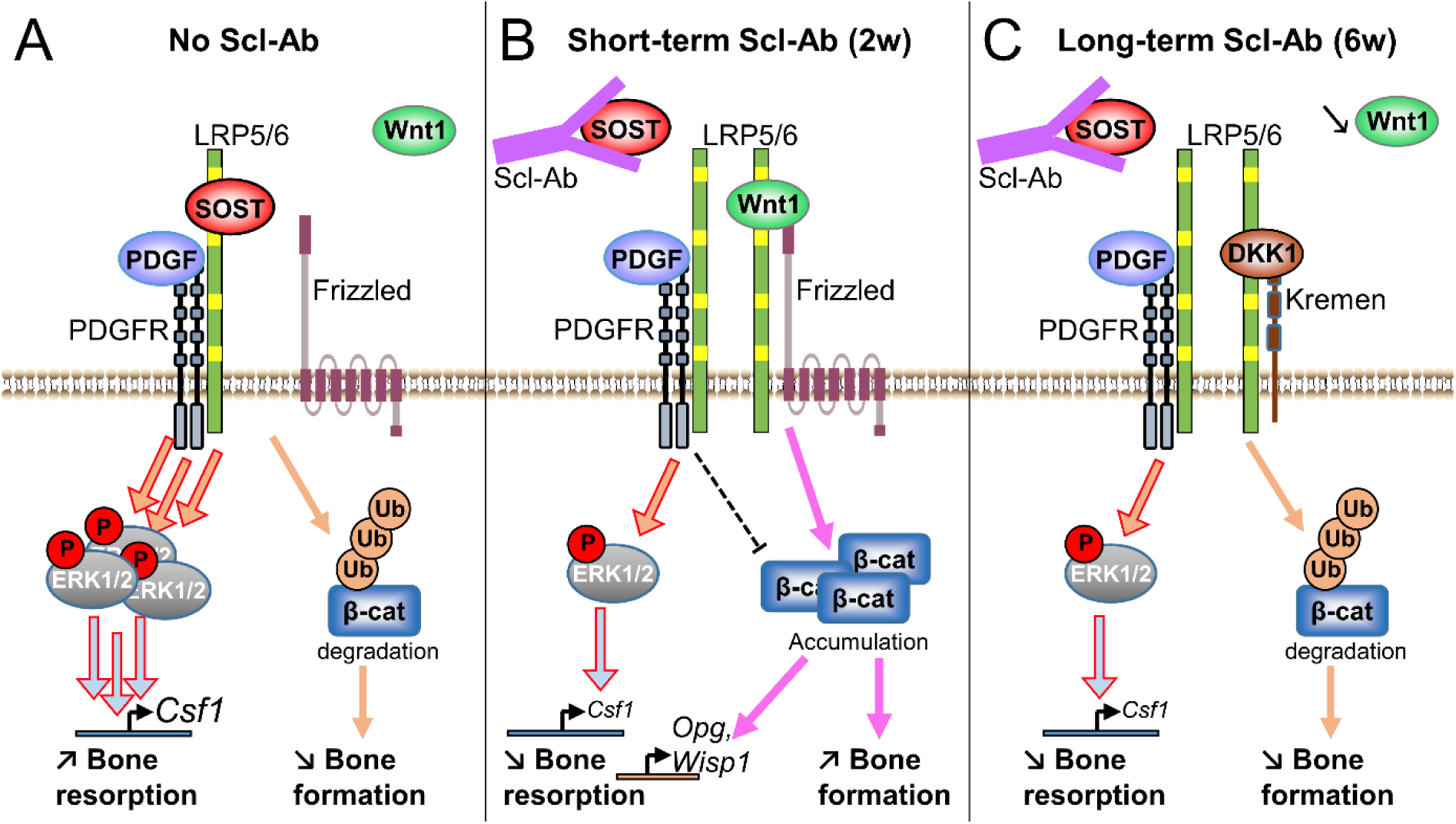
Proposed molecular mechanisms for Scl-Ab actions. (A) Besides its function as a Wnt/LRP6 antagonist that promotes β-catenin degradation by the proteasome, sclerostin forms a ternary complex with LRP6 and PDGFRs, leading to co-activation of PDGF-BB/PDGFRs/ERK1/2 signaling and *Csf1* expression. (B) Short-term Scl-Ab exposure (2 weeks) prevents sclerostin binding to LRP6, thereby promoting Wnt1 class-induced β-catenin accumulation and signaling, and preventing sclerostin-mediated co-activation of PDGFRs/ERK1/2 signaling and *Csf1* up-regulation. In this context, residual PDGFR signaling inhibits Wnt/β-catenin signaling. (C) During prolonged Scl-Ab exposure (6 weeks), a negative feedback mechanism, consisting in elevated expressions of the Wnt signaling inhibitor DKK1 and decreased expressions of Wnt1 class of ligands, attenuates Wnt/β-catenin signaling, while Scl-Ab still continues to prevent sclerostin-mediated co-activation of osteocatabolic PDGF-BB/PDGFRs/ERK1/2 signaling.

Scl-Ab treatment exerted sustained inhibition of bone resorption in control mice, as observed in humans (7–10). Scl-Ab administration transiently enhanced expression of *Wisp1*, a Wnt1-responsive negative regulator of osteoclastogenesis (22), and continuously reduced expression of *Csf1*, encoding an essential growth factor for osteoclastogenesis (Figure 3) (14). The differential regulation between *Wisp1* and *Csf1* expressions by Scl-Ab suggested that sclerostin could regulate *Csf1* expression independently of Wnt/β-catenin signaling inhibition. Interestingly, Scl-Ab treatment could not further reduce *Csf1* expression in *Pdgfr cKO* mice (Figure 3), thus indicating that PDGFR signaling is sufficient and necessary to explain sclerostin effects on *Csf1* expression and bone resorption. At the cellular level, short-term Scl-Ab treatment induced a slight additional decrease in osteoclast surfaces while prolonged Scl-Ab exposure did not further reduce them in *Pdgfr cKO* mice (Figure 3), thus reflecting the combination of a transient stimulatory effect on Wnt/β-catenin-mediated *Wisp1* and possibly *Opg* expressions, and a continuous inhibitory effect on PDGFR-mediated *Csf1* expression (Figure 7). Altogether, those findings provide the cellular and molecular mechanisms involved in the biphasic inhibitory effects of Scl-Ab on bone resorption observed in clinical trials, i.e., a sharp decrease in serum CTX levels at 1 month, a return close to baseline at 3 months, followed by a progressive and continuous reduction at 6 and 12 months (7–9).

In line with the continuous inhibitory effect of Scl-Ab on PDGFR-mediated *Csf1* expression, prolonged anti-catabolic effect of Scl-Ab was associated with diminished expressions of PDGFR target genes *c-Myc* and *Ccl2* in bone (Figure 4). The fact that Scl-Ab could prevent sclerostin-mediated potentiation of PDGF-BB-dependent PDGFRs activation and *Csf1* expression in pre-osteoblast cultures, and calcitriol-induced osteoclast formation and activity in pre-osteoblast/osteoclast co-cultures also supports our in vivo observations (Figure 5). Most importantly, we demonstrated that sclerostin, in contrast to DKK1, could function as a co-activator of PDGFRs and downstream ERK1/2 signaling pathways, independently of Wnt1 and Wnt/β-catenin signaling inhibitors (Figure 6), thereby explaining why bone resorption remains inhibited by prolonged Scl-Ab treatment despite the downregulation of bone formation (Figure 7). At a molecular level, heterodimerization between PDGFRs and LRP6 likely contributes to sclerostin effects on PDGFR signaling in pre-osteoblasts (Figures 5 and 7).

In conclusion, we identified a new pathway for sclerostin effects on bone resorption and provide an explanation for the dissociation of Scl-Ab long-term therapeutic efficacy on bone resorption versus formation. Indeed, Scl-Ab anti-catabolic effects on the skeleton occur by repressing PDGFR signaling and M-CSF expression in osteoblast lineage cells in a Wnt-independent manner. In addition, although it remains to be formally shown that activating mutations of PDGFRβ in osteoprogenitors contributes to osteopenia and occurrence of fractures in patients with Penttinen syndrome or Kosaki overgrowth syndrome (23), Scl-Ab could be useful to treat those pathological conditions. Finally, our findings also suggest that combination of PDGFR inhibition and sclerostin neutralization could represent a powerful approach to rapidly increase bone mass and strength in patients with osteolytic lesions provoked by multiple myeloma or bone metastases involving excessive PDGFR activity (24, 25).

## Materials and Methods

### Mice and Scl-Ab treatment

*Pdgfr cKO* mice in which *Pdgfra* and *Pdgfrb* genes can be selectively deleted under the control of an *Osterix* promoter following cessation of doxycycline treatment (Tet-Off system) were generated as previously described (17). Since both PDGFRs can compensate for the loss of the other and exert redundant function in osteoblast lineage cells (17), we used *Pdgfr cKO* mice in our experiments. *Osx-Cre* mice were used as control animals. All mice were on a C57BL/6J genetic background. 16-week-old male *Osx-Cre* and *Pdgfr cKO* mice were randomly assigned to receive subcutaneous injections of 25 mg/kg Scl-Ab (r13c7, provided by UCB Pharma and Amgen Inc.) or an equivalent volume of saline solution twice a week for 2 weeks (bone formation rate peaked after 2 weeks of Scl-Ab treatment) or 6 weeks (bone formation rate returned to basal level after 6 weeks of Scl-Ab treatment) (Figure 1A) (21). *Cre* expression and consequent *Pdgfra* and *Pdgfrb* inactivation were induced one week prior to the onset of Scl-Ab treatments by stopping doxycycline administration (Figure 1A). Mice (3 to 6 animals per cage) were maintained under standard non-barrier conditions, exposed to a 12-hour light/12-hour dark cycle and had access to mouse diet RM3 containing 1.24 % calcium and 0.56 % available phosphorus (SDS, Betchworth, UK) and water *ad libitum*. Experimental units were single animals. Mice treatments (Veh or Scl-Ab) and endpoint measurements (by µCT and histomorphometry) were performed by different investigators. Investigators were blinded during endpoint measurements.

### Bone phenotyping

Mice were sacrificed and their bones were excised for µCT analyses. Trabecular bone microarchitecture of proximal tibiae (100 slices from the beginning of secondary spongiosa), and cortical bone geometry of tibial midshafts (50 slices) were assessed using µCT (Viva-CT40, Scanco Medical, Switzerland) employing a 12-μm isotropic voxel size.

To measure dynamic indices of bone formation, mice received subcutaneous injections of calcein (10 mg/kg body weight; Sigma) at 9 and 2 days before euthanasia. Formalin-fixed undecalcified femurs were embedded in methylmethacrylate (Merck). 8-µm transversal sections of midshafts and 8-µm sagittal sections of distal femurs were cut and mounted unstained for fluorescence visualization. Additional sagittal sections were stained with Goldner trichrome for osteoblast counting or with tartrate-resistant acid phosphatase (TRAP) substrate for osteoclast counting. Histomorphometric measurements were carried out using a Nikon Eclipse microscope and the BioQuant software.

### Biochemistry

Serum levels of PINP and TRAcP 5b were determined by using immunoassay kits (Immunodiagnostic Systems Ltd). M-CSF protein levels were determined in bone marrow supernatants (obtained by centrifugation at 1000 g for 10 minutes) or in cell culture medium using Quantikine ELISA kits (R&D Systems). Results obtained with bone marrow supernatants were normalized to total protein levels.

### Osteoblast cultures

Primary pre-osteoblasts were isolated from long bones of *Pdgfra^f/f^;Pdgfrb^f/f^* (*Pdgfr f/f*) mice as previously described (26). Briefly, bone chips were prepared from cleaned long bones and digested in 1 mg/mL collagenase II (Sigma) for 90 minutes at 37 °C. Bone pieces were washed several times and incubated in α-MEM (Amimed, Bioconcept) containing 10% FBS (Gibco) for 9 days to allow cell migration from bone fragments. At that point, cells and bone chips were trypsinized (with trypsin/EDTA from Sigma) and passaged at a split ratio of 1:3. At the second passage, bone chips were removed. Medium was changed every 2-3 days. Pre-osteoblasts at passages 3-4 were used for in vitro experiments. *Pdgfr f/f* pre-osteoblasts were infected with 400 moi of empty or Cre-expressing adenoviruses (Vector Biolabs) to obtain *Pdgfr f/f* (control) and *Pdgfr cKO* pre-osteoblasts. Osteoblast differentiation was determined by incubating confluent pre-osteoblast cultures in osteogenic medium containing α-MEM, 10% FBS, 0.05 mM L-ascorbate-2-phosphate (Sigma) and 10 mM β-glycerphosphate (AppliChem GmbH) in the presence of vehicle (Veh), 100 ng/mL Wnt1-sFRP1 (R&D Systems) ± 500 ng/mL recombinant sclerostin (Peprotech).

### Co-cultures

For co-culture experiments, primary *Pdgfr f/f* pre-osteoblasts infected with 400 moi of empty or Cre-expressing adenoviruses were seeded at 30000 cells per well in 24-well plates. The day after, non-adherent bone marrow cells isolated from wildtype mice were seeded over pre-osteoblasts at 300000 cells per well in α-MEM supplemented with 10% FBS and treated with Veh, 10^-8^ M 1,25-dihydroxyvitamin D3 (calcitriol, Vit.D3), or/and 250 ng/mL recombinant murine sclerostin (Peprotech), with or without 1.25 µg/mL Scl-Ab or 500 ng/mL anti-M-CSF antibody (#AF416 from R&D Systems). After 8 days, co-cultures were fixed and stained, and multinucleated TRAP-positive cells were counted. To evaluate in vitro osteoclast-mediated demineralization, co-cultures were performed under the same conditions in Corning® Osteo Assay Surface multiple well plates for 15 days. Multiple well plates were cleaned with a bleaching solution and observed under an inverted phase-contrast microscope (Nikon Eclipse TE2000). Synthetic matrix demineralization was quantified using Image J software.

### Immunoprecipitations

Confluent pre-osteoblast cultures were pre-treated with Veh or 500 ng/mL recombinant sclerostin for 2 hours, and then treated with 25 ng/mL PDGF-BB for 15 minutes. Cell lysates were prepared by incubating osteoblast cultures in lysis buffer containing 1% NP-40 and phosphatase/protease inhibitors at 4 °C for 30 minutes. Cells were then scraped and sonicated on ice for 10 seconds. Lysates were then centrifuged at 6000g for 30 minutes at 4 °C. Cell lysates were incubated with 1:200 anti-LRP6 antibody (#3395 from Cell Signaling) or isotype control overnight at 4 °C. The day after, immunocomplexes were incubated with pre-washed Protein G Magnetic beads (#70024 from Cell Signaling) under agitation for 1 hour at room temperature. Then, beads were pelleted by using a magnetic separation rack and washed 3 times with cell lysis buffer. Immunocomplexes attached to beads were diluted with equal volumes of 2-fold concentrated loading buffer and heated at 70 °C for 30 minutes. Immunocomplexes were separated from magnetic beads by using a magnetic separation rack and collecting supernatants. Finally, supernatants were subjected to western blot analyses as described below.

### Western Blots

To measure PDGFR signaling, confluent pre-osteoblast cultures were pre-treated with Veh or 500 ng/mL recombinant sclerostin ± 2.5 µg/mL Scl-Ab for 1 hour, and then treated with 15 ng/mL PDGF-BB for 15 minutes. Pre-osteoblast cultures were rapidly frozen in liquid nitrogen and stored at -80 °C until their use for analysis. Cell lysates were prepared by incubating cell cultures in RIPA buffer containing phosphatase and protease inhibitors at 4 °C for 30 minutes. Lysates were then centrifuged at 6000g for 30 minutes. Lysate supernatants were diluted with equal volumes of 2-fold concentrated reducing sample buffer. Those mixtures were then heated at 70 °C for 30 minutes and subjected to gel electrophoresis on 6% to 15% gels. Proteins were electro-transferred to Immobilon P membranes and immunoblotted with specific antibodies: anti-PDGFRα (#3174), anti-PDGFRβ (#3169), anti-p-PDGFR (#3170), anti-LRP6 (#3395), anti-p-LRP6 (#2568), anti-p-STAT3 (#9138), anti-p-SHP2 (#3751), anti-p-ERK1/2 (#9106), anti-ERK1/2 (#4695), anti-p-β-catenin (Ser552) (#9566), anti-pan-Actin (#8456) (from Cell Signaling), and anti-M-CSF (#AF416 from R&D Systems). Detection was performed using peroxidase-coupled secondary antibody, enhanced chemiluminescence reaction, and visualization by using G:Box gel analysis system (Syngene). Reprobed membranes were stripped according to the manufacturer’s protocol.

### RNA isolation and real-time PCR

Total RNA was extracted from tibial metaphyses (whose bone marrow was removed by centrifugation at 16200g for 20 seconds) or primary pre-osteoblast cultures using Tri Reagent^®^ (Molecular Research Center) and purified using a RNeasy Mini Kit (Qiagen). Single-stranded cDNA was synthesized from 2 µg of total RNA using a High-Capacity cDNA Archive Kit (Applied Biosystems) according to the manufacturer’s instructions. Real-time PCR was performed to measure the relative mRNA levels using the QuantStudio™ 5 Real-Time PCR System with SYBR Green Master Mix (Applied Biosystems). The primer sequences are described in supplementary table 1. Melting curve analyses performed at the completion of PCR amplifications revealed a single dissociation peak for each primer pair. The mean mRNA levels were calculated from triplicate analyses of each sample. Obtained mRNA level for a gene of interest was normalized to β2-microglobulin mRNA level in the same sample.

### Statistics

A sample size of 5 mice/group was required in order to detect a difference of 30% in fractional trabecular osteoclast surface (SD=30%) between groups at the significance level of 0.01 and a power of 80%. In vitro experiments were performed in triplicate and independently repeated 3 or 4 times. Interactions between effects of treatments and those of genotypes, were analyzed by using 2-way ANOVA followed by Tukey post hoc tests. Interactions between effects of treatments, those of treatment durations and those of genotypes, were analyzed by using linear mixed-effects models followed by Tukey post hoc tests.

## Study approval

All performed experiments were in compliance with the guiding principles of the *Guide for the Care and Use of Laboratory Animals* (8^th^ edition) and approved by the Ethical Committee of the University of Geneva School of Medicine and the State of Geneva Veterinarian Office.

## Data availability

All data supporting the findings of this study are reported in the Supporting Data Values file and available from Zenodo (https://zenodo.org) with the identifier doi:10.5281/zenodo.7670092.

## Author Contributions

C.T.: Study design; methodology; data acquisition; data analysis; funding acquisition; investigation; project administration; supervision; validation; writing-original draft; writing-review and editing. P.A.: methodology; data acquisition; writing-review and editing. J.B.: methodology; data acquisition; writing-review and editing. J.C.: validation; writing-review and editing. S.F.: funding acquisition; validation; writing-review and editing.

## Acknowledgments

We would like to thank Dr. Gill Holdsworth from UCB Pharma for supplying Scl-Ab and the careful reading of the manuscript. This work was supported by the Swiss National Science Foundation (310030_185230), by the Fondation pour la Recherche sur l’Ostéoporose et les Maladies Osseuses, and by the Novartis Foundation for medical-biological research (Project number 16C212 - Basel, Switzerland).

