## Supplemental Materials for "Sclerostin blockade inhibits bone resorption through PDGF receptor signaling in osteoblast lineage cells"

**Supplemental Table 1**. List of primers used in real-time PCR.

| **Gene** | **Primer** | **Sequence** |
|---|---|---|
| *B2m* | F | CACTGACCGGCCTGTATGCT |
| | R | GTATGTTCGGCTTCCCATTCTC |
| *Rankl* | F | GCACACCTCACCATCAATGC |
| | R | AGCCTCGATCGTGGTACCAA |
| *Opg* | F | GACAACGTGTGTTCCGGAAA |
| | R | GGTAGGAACAGCAAACCTGAAGA |
| *Csf1* | F | CAGCTGCTGGAGAAGATCAAGA |
| | R | CCACATCTCGGCTAGAGCACTT |
| *Twist1* | F | GGCCAGGTACATCGACTTCCT |
| | R | CGCTCGTGGGCCACATAG |
| *Wisp1* | F | GGGCCTCTACTGCGATTACAGT |
| | R | CCACCTGTGCACACACTCCTA |
| *Sost* | F | AGACCTCCCCACCATCCCTAT |
| | R | TGTCAGGAAGCGGGTGTAGTG |
| *Dkk1* | F | CAACGCGATCAAGAACCTG |
| | R | TCCCGCCCTCATAGAGAAC |
| *Wnt1* | F | GCAAATGGCAATTCCGAAA |
| | R | GGAGGTGATTGCGAAGATGAA |
| *Wnt10b* | F | AATGCGGATCCACAACAACA |
| | R | GCACTTCCGCTTCAGGTTTT |
| *Pgdfra* | F | CAACCACACTCAGACGGATGA |
| | R | GGCCATGTCTGGGTCTGGTA |
| *Pdgfrb* | F | CTGAGTGATTCGGGCACCTATAC |
| | R | ACGTAGCCATTCTCGATCACAGA |
| *Pdgfb* | F | GGTCCAGGTGAGAAAGATTGAGA |
| | R | GGTGGTCCTCCAAGGTCACT |
| *Myc* | F | GCCCCTAGTGCTGCATGAG |
| | R | TCCACAGACACCACATCAATTTC |
| *Ccl2* | F | GGCTCAGCCAGATGCAGTTAA |
| | R | GCCTACTCATTGGGATCATCTTG |
| *Runx2* | F | CGGACGAGGCAAGAGTTTCA |
| | R | GGGACCGTCCACTGTCACTT |
| *Col1a1* | F | CTGGCCTTGGAGGAAACTTT |
| | R | GCACGGAAACTCCAGCTGAT |
| *Alpl* | F | AGATGGCCTGGATCTCATCAGT |
| | R | GTTCAGTGCGGTTCCAGACATA |
| *Ocn* | F | GGAGGGCAATAAGGTAGTGAACAG |
| | R | CACAAGCAGGGTTAAGCTCACA |

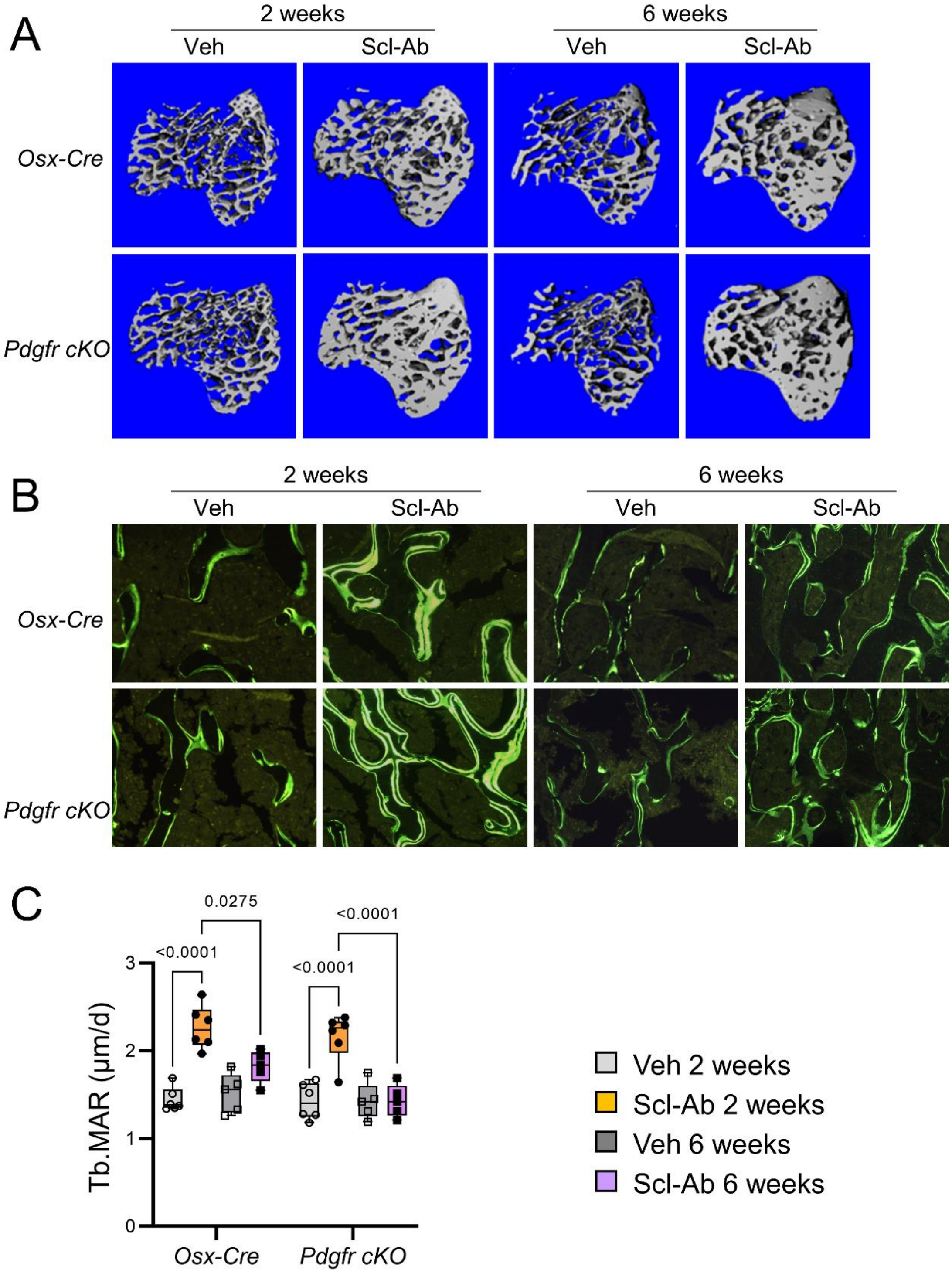


**Supplemental Figure 1. Effects of Scl-Ab treatment on trabecular bone mass and bone formation in *Osx-Cre* and *Pdgfr cKO* mice.** (A) Representative reconstructed µCT images of proximal tibiae from 4-month-old *Osx-Cre* and *Pdgfr cKO* (*Pdgfra cKO;Pdgfrb cKO*) male mice that received subcutaneous injections of saline solution (Veh) or 25 mg/kg Scl-Ab twice a week for 2 or 6 weeks. *Cre* expression or/and conditional gene deletion were induced (by stopping doxycycline treatment) one week prior the beginning of Scl-Ab treatment. (B) Representative images of calcein-labeled histological sections of distal femurs. (C) Trabecular mineral apposition rate (Tb.MAR) measured at the secondary spongiosa of distal femurs (n=5-

6 per group). Interactions between effects of genotypes, those of treatments, and those of treatment durations were analyzed by linear mixed-effects models. Differences between genotypes, between treatments or between durations were analyzed using Tukey post hoc tests.

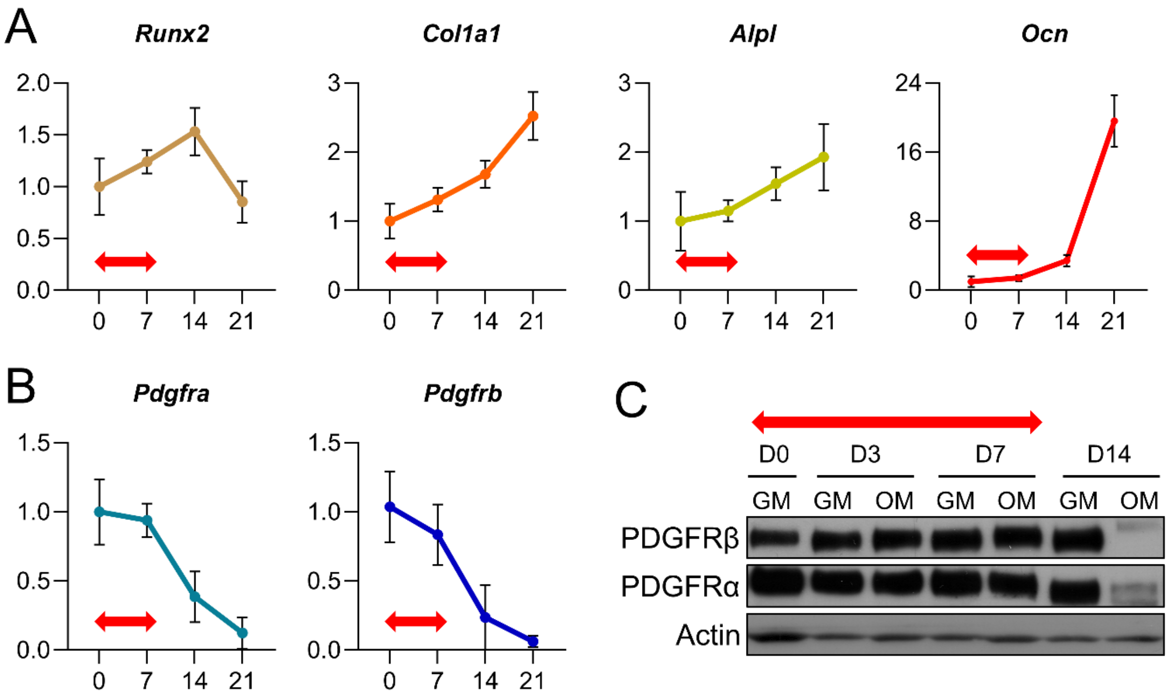


**Supplemental Figure 2. Expression of PDGFRα/β in osteoblast lineage cells.** (A) Quantitative RT-PCR analyses of expressions of *Runx2*, *Col1a1*, *Alpl* and *Ocn* in primary osteoblast lineage cells cultured for 0, 7, 14, and 21 days after confluence in osteogenic medium. (B) Quantitative RT-PCR analyses of expressions of *Pdgfra* and *Pdgfrb* in primary osteoblast lineage cells cultured for 0, 7, 14, and 21 days after confluence in osteogenic medium. (C) Western blot analyses of PDGFRα and PDGFRβ levels in primary osteoblast lineage cells cultured for 0, 3, 7, and 14 days after confluence in growth medium (GM) or osteogenic medium (OM). Red arrows indicate the differentiation stage of osteoblast lineage cells (i.e., pre-osteoblasts) used in cell culture experiments.

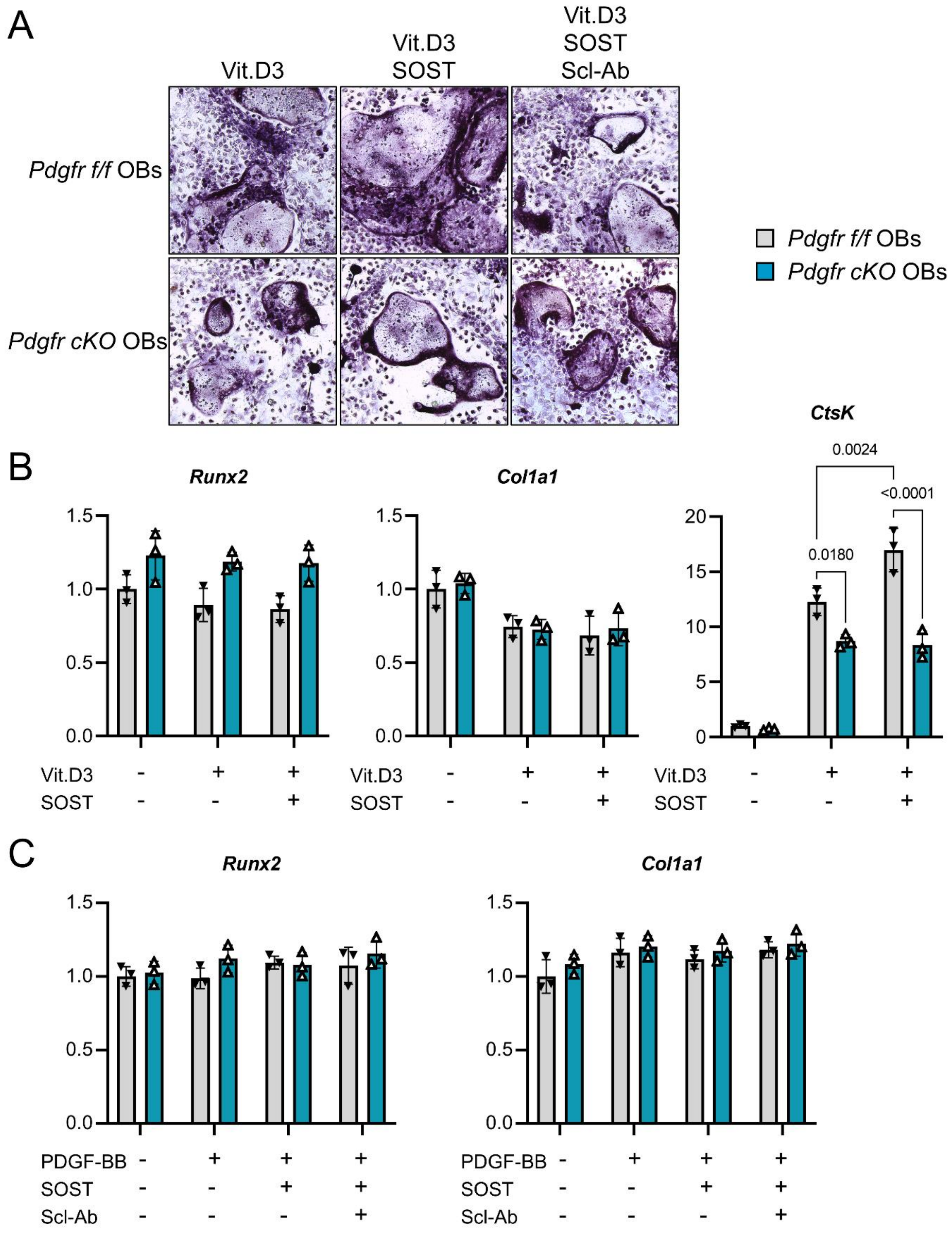


**Supplemental Figure 3. Effects of sclerostin and Scl-Ab on osteoblastogenesis and calcitriol-induced osteoclastogenesis in vitro.** (A) *Pdgfr*$^{f/f}$ (control) and *Pdgfr cKO* (without PDGFRs) osteoblasts were co-cultured with non-adherent bone marrow cells isolated from wildtype mice in the presence of Veh, $10^{-8}$ M 1,25-dihydroxyvitamin D3 (calcitriol, Vit.D3), or/and 250 ng/mL recombinant sclerostin (SOST), with or without 1.25 µg/mL Scl-Ab during 8 days before quantification of TRAP-positive multinucleated cells. (B) Measurements of *Runx2*, *Col1a1* and *CtsK* expression by quantitative RT-PCR in the same co-cultures. (C)

*Pdgfr$^{f/f}$* and *Pdgfr cKO* osteoblasts were pre-treated with Veh, 250 ng/mL SOST ± 1.25 µg/mL Scl-Ab for 1 hour, and treated with Veh or 25 ng/mL PDGF-BB for 24 hours before measurements of *Runx2* and *Col1a1* expression by quantitative RT-PCR.
